# motifNet: A Neural Network Approach for Learning Functional Sequence Patterns in mRNA

**DOI:** 10.1101/2022.12.20.521305

**Authors:** Kaifeng Deng, Zhengchang Li, Wenqing Wei, Yang Liu

## Abstract

We present a new approach for predicting functional sequence patterns in mRNA, known as motifs. These motifs play an important role in understanding the mechanisms of the cell life cycle in clinical research and drug discovery. However, many existing neural network models for mRNA event prediction only take the sequence as input, and do not consider the positional information of the sequence. In contrast, motifNet is a lightweight neural network that uses both the sequence and its positional information as input. This allows for the implicit neural representation of the various motif interaction patterns in human mRNA sequences. The model can then be used to interactively generate motif patterns and the positional effect score in mRNA activities. Additionally, motifNet can identify violations of motif patterns in real human mRNA variants that are associated with disease-related cell dysfunction.

## 1 Introduction

Functional sequence patterns in messenger RNA (mRNA) play a critical role in regulating mRNA expression, stability, localization, translation initiation, and cellular function. It is important to build analysis tools to predict those mRNA events. In recent years, deep learning models have made significant progress in modeling biological data. In the realm of mRNA sequences, however, researchers have largely relied on one-hot encoding of the four nucleotides (A, U, C, and G) in the sequence as a data representation method. This explicit representation of mRNA sequences has limitations when it comes to modeling and feature interpretation. For example, a pair of motifs with the same sequence pattern will be encoded equally in most network models, despite the fact that different positional patterns within the mRNA sequence can have a dramatic impact on cellular function.

For example, in the genetic code, the sequence “AUG” is known as the start codon, as it signals the start of protein translation. The start codon is typically located in the 5’ untranslated region (UTR) of mRNA, which is the region of the mRNA molecule that is not translated into protein. The position of the start codon relative to the reading frame of the mRNA sequence determines whether it is “in-frame” or “out-of-frame”. An in-frame start codon is one that is in the correct reading frame, such that the subsequent nucleotides are translated into the correct sequence of amino acids. In contrast, an out-of-frame start codon is one that is not in the correct reading frame, resulting in a shift in the sequence of amino acids and potentially causing the translation to stop prematurely. The difference between in-frame and out-of-frame start codons can have a significant impact on the final protein product, as the sequence of amino acids determines the protein’s structure and function.

When encoding mRNA sequences using sequence models, in-frame and out-of-frame AUG codons will be encoded as the same feature, as they have the same sequence of nucleotides. However, the biological activity of these two codons can be quite different, as the position of the start codon relative to the reading frame can impact the final protein product. An in-frame AUG codon will be translated into the correct sequence of amino acids, resulting in a functional protein. In contrast, an out-of-frame AUG codon may be translated into a truncated protein with reduced or altered function. This difference in biological activity can have significant consequences, as the function of a protein can be crucial for cellular processes and overall health. By using sequence models that encode in-frame and out-of-frame AUG codons as the same feature, the positional information that is critical for determining their biological activity is lost. This can result in inaccurate predictions and a lack of interpretability in the model. In order to accurately model the biological activity of mRNA sequences, it may be necessary to use alternative methods that can capture the positional information of functional elements in the sequence.

Recent attempts have been made to address these limitations. Jaganathan et al. (2019) developed a 32-layer ResNet model called spliceAI for accurate splicing event prediction. In order to consider the impact of distal sequences in mRNA, the spliceAI model takes a 15,000 nucleotide-long mRNA sequence as input and does not have any down-sampling in all layers. This makes the model very large and time-consuming to train, and the final model is not interpretable, making it difficult to understand how alternative splicing events occur. Instead, the authors used a mutagenesis method to alter each input sequence nucleotide to see the delta change in the final splicing event probability.

The use of Implicit Neural Representations (INR) offers an alternative way to parameterize features in mRNA sequences. INR has been widely used in image and 3D shape representation, such as Neural Radiance Fields (NERF) (Mildenhall et al., 2020). INR is also known as a coordinate-based representation, as the model inputs are coordinate positions rather than the original discrete data such as pixels or audio signals (Rougier, 2009). Implicit representation modules map data positions to target labels at the coordinate using a continuous function. Implicit representation learning has the natural benefit of characterizing positional and interval information in data. For example, Sitzmann et al. (2020) found that 3D shapes with many edges and circles are easier to learn with INR. In the case of biological sequences, positional features are even more common, such as the amino acid coding codon in RNA and tandem repeats in DNA. Sequence mutation resulting in an increase of tandem repeats can increase the risk of ASD disease (Mitra et al., 2021; Trost et al., 2020). RNA functional elements are even more sensitive to global and local position. For example, in the 5’UTR region of mRNA, an in-frame upstream AUG compared to an out-of-frame AUG can have a dramatic impact on the final translation, even if there is only a single nucleotide shift.

Another benefit of using INR in mRNA is reducing memory requirements for extremely long sequences. By learning a function over sequence space, motif encoder layers can focus only on important regions rather than scan the whole sequence. Thus the method we propose here provides an approach to detect new function regulated sequence motifs in mRNA.

In summary, our main contributions include:

- Defining a generalized framework for position-specific sequence pattern extraction and interpretation using INR in mRNA sequence function mapping.
- Demonstrating an approach to evaluate the functional effects of motif patterns in mRNA sequences, including the exploration of motif interaction patterns and their potential correlation with disease variants.
- Developing a new technique for reducing memory requirements in splicing data analysis by using positional activation modules in lighter networks

## 2 Related Works

### 2.1 Implicit neural representations in different fields

Recent studies have shown the effectiveness of using convolution or fully connected networks as implicit representations for 2D images (Shaham et al., 2021; Chan et al., 2020) or 3D scenes (Mildenhall et al., 2020; Mescheder et al., 2019; Niemeyer & Geiger, 2020). These models are typically trained from a single image or synthetic 3D data. Tissieres et al. (2019) model EEG and fMRI signal with INR to learn position-linked feature. However, implicit representation in the field of biological sequences has not been explored yet, due to the difference of data distribution.

### 2.2 Biological motifs extraction by neural network

Biological motif extraction by neural networks refers to the use of machine learning techniques, specifically neural networks, to identify patterns in biological sequences such as DNA or RNA. These patterns, known as motifs, can be indicative of specific functions or properties of the sequence, such as binding to a particular protein or being involved in a particular biological process.

One example of such a method is PrismNet, which was developed by Sun et al. (2021) to predict RNA binding protein (RBP) motifs by training on RBP crosslinking immunoprecipitation sequencing (CLIP-seq) datasets. The attention mechanism is used in PrismNet to improve the prediction of motifs. Other methods for identifying motifs in biological sequences include using convolutional networks or evaluating the effects of specific motifs through frequency statistics or saturated mutagenesis. Sample et al. (2019) used a 3-layers convolution network for translation prediction, but the only way to visualize functional elements is by sequence frequency statistics from different translation levels. Linder et al. (2022) used scrambler network to calculate the important mask of 5’UTR sequence, yet they didn’t solve the problem of dynamic length and continuity of motif. SpliceAI evaluates the effects of branch point TACTAAC to acceptor sites by continuous mutation of this motif in every position of the whole sequence. The common limitation of these methods is that if we don’t have the domain knowledge of a certain motif pattern, there is no way to search a sequence pattern along the sequence samples at the first stage. Koo & Ploenzke (2020) made a further improvement to genomic sequence motifs representation by adding exponential activations to convolutional networks. In summary, these methods often relied on manual feature engineering and required prior knowledge of the motif pattern being searched for.

### 2.3 Positional interacted motif patterns discovered by neural networks

Avsec et al. (2019) explored transcription factor (TF) motifs and discovered a 10.5 bp period interaction pattern in DNA sequences using a convolution network called BPnet. However, their approach required specifying TF motif patterns for interaction search.

## 3 Method

Generally, we are representing sequence information in a form of:

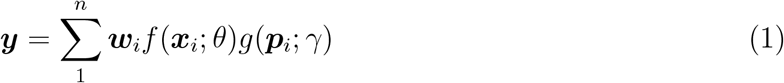

Where ***y*** denotes the mRNA prediction tasks like translation efficiency or splicing events, *f* (***x***_*i*_; *θ*) denotes a sequence pattern filtering function for motif extraction from sequences ***x***_*i*_. ***p***_*i*_ denotes positional information of the motif. The position encoding module of motifNet is a learnable formulation parameterized by *γ* which takes the RNA sequence coordinates ***p***_*i*_ as input and output a suitable positional effect score evaluating the function regulation. ***w***_*i*_ is the contribution weight matrix to the final output y value.

In practice we build two neural network encoder for positional and structural information understanding, as depicted in Fig. 1.

**Figure 1:**
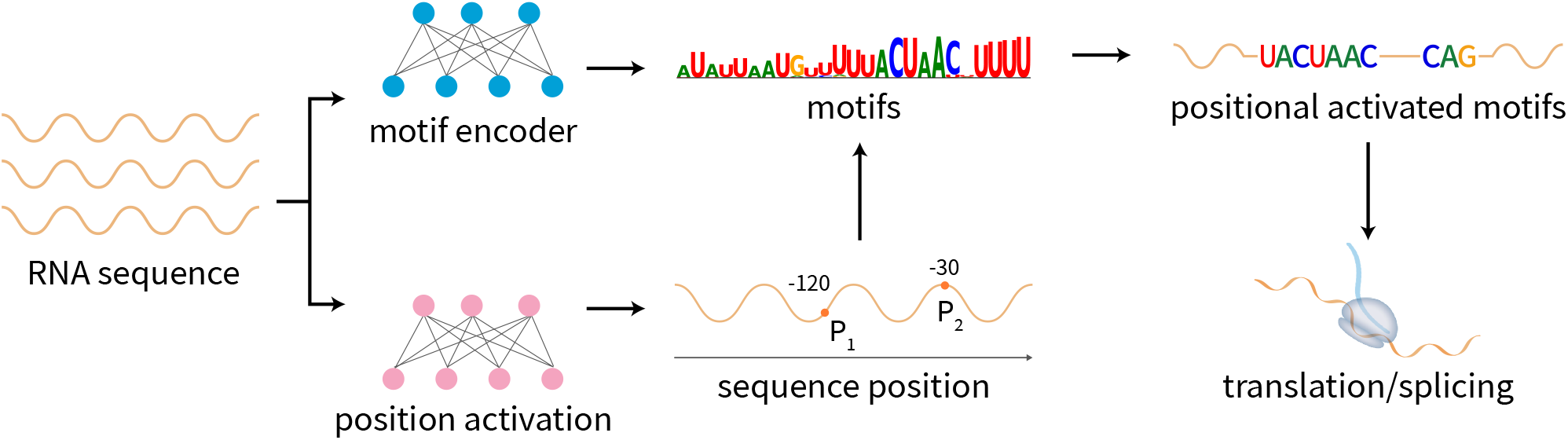
Overview of motifNet workflow. RNA sequences were numerically encoded and fed into motif encoder while normalized samples coordinates were used for implicit representation function optimization. Then the positional activated motifs were multiplied by a interaction weight matrix for final translation or splicing prediction.

### 3.1 Motifs pattern representation

In order to visualize the optimized pattern parameters in *f* (***x***_*i*_; *θ*), we transform weight matrix *θ* into a Position-weight-matrices(PWM). PWM is also known as position-specific weight matrix(PSWM), which is a traditional approach to represent biological sequence patterns by calculation of occurrence frequency of the elements in sequences, which is A,U,C,G bases specifically for mRNA. Here the position importance of sequence pattern was produced by 1D convolution where kernel weight was used as a similar metric to represent the recognized motif patterns. For a sequence *S*, the probability, or pattern activation level that it contained a motif pattern ***a*** is defined as:

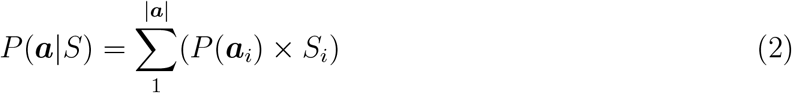

For each nucleotide base *S*_*i*_ it will multiply the weight score *P* (***a***_*i*_) at its position. Similar motif patterns in sequence will be activated as depicted in Fig. 2. PWM is also able to encode positional dependency as long as we lengthen the kernel in 1D convolution, or add multiple layers to enlarge the receptive field. However, larger kernel size or deeper network is infeasible due to the memory limitation of hardware. Receptive field size *r*_0_ of the network can be computed as 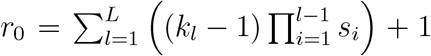. If all strides are 1, as implemented in spliceAI, then the receptive field will simply be the sum of kernel shape (*k*_*l*_ − 1) over all layers plus 1 (Araujo et al., 2019). If we use *k*_*l*_ = 3 then we need a 2500 layers of convolution to cover the whole sequence! Even if we enlarge the kernel size to 11, reducing the number of layers to 250, it is still two huge to build. Thus we proposed position encoder *g*(***p***_*i*_; *γ*) and down-sampling network *D*(***x***_*i*_) to improve model long term pattern recognition with memory-efficient. This part will be explained in detail in the following sections.

**Figure 2:**
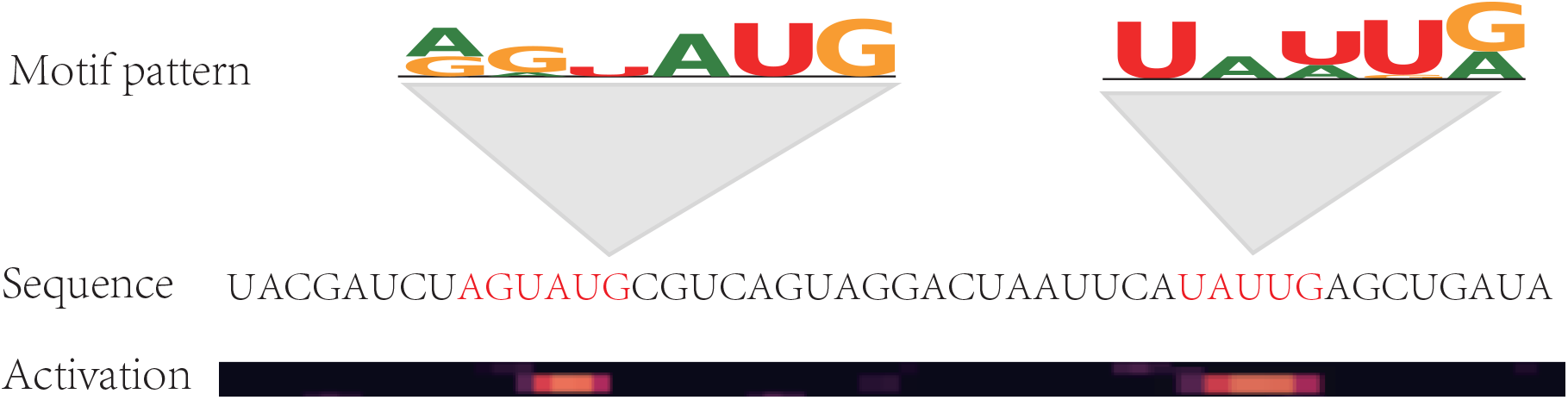
Motifs activation were calculated by matrix multiplication of one hot encoded sequence pattern and PWM.

### 3.2 Combining shallow and deep layers for motifs extraction

Here we proposed a framework to integrate deep network feature into a shallow layer for better motif interpretation. As defined in Equation (1), motif pattern extraction is done by network module *f* (***x***_*i*_; *θ*), which can be further divided into two parts: a PWM-like pattern activation layer *C*(***x***_*i*_) and a deep multi-layers down-sampling network *D*(***x***_*i*_).

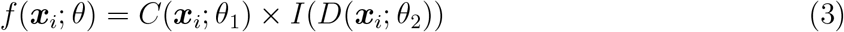

In practice we used a convolutional neural network (CNN) model to extract features from mRNA sequences. In this equation, *f* (***x***_*i*_; *θ*) represents the combined feature activation, which captures both the direct motif patterns learned by the first layer of the CNN model and deeper features learned by the model from the deeper layers. *C*(***x***_*i*_; *θ*_1_) represents the first layer of the CNN model, which is parameterized by the weights *θ*_1_. This layer can be used as a position weight matrix (PWM) representation of the motif patterns learned by the model. *D*(***x***_*i*_; *θ*_2_) is a multi-layer feature extractor module of the CNN model, which is parameterized by the weights *θ*_2_. The output of this module, denoted as *D*(***x***_*i*_), is a feature representation of the input sequence that is shorter in length than the original sequence. The linear interpolation function, denoted as *I*, is used to map the feature representation back to a mask with the same length as the original sequence. This mask, referred to as the activation mask, is used to identify which positions in the mRNA sequence are most important for the model’s predictions.

This allows us to visualize the different motif patterns learned by the model at different positions in the mRNA sequence. In addition to the direct motif patterns learned by the first layer of the CNN model, which we refer to as the baseline motifs, we also consider the combined feature activation, denoted as *f* (***x***_*i*_; *θ*), which we refer to as the motifNet activated motifs. These motifs capture both the direct motif patterns from the first layer of the CNN model and context features learned by the model from the deeper layers.

### 3.3 Position effect score evaluation of a motif

Following our motif extraction definition above in Equation (3), we next adding positional variation in *D*, which means that we are giving both the original sequence ***x***_*ori*_ and alternative variant ***x***_*alt*_ to the model as *C*(***x***_*ori*_, *θ*_1_) × *I*(*D*(***x***_*alt*_) with positional input *g*(***p***_*alt*_; *γ*). Now by observing the probabilities change of final output *y*, we can get the sequence wide motif effect in different position. The overall optimizing method can be written as:

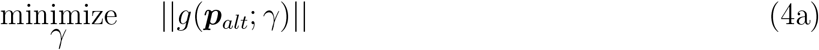

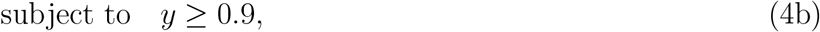

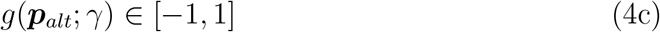

We iterate user given motif pattern ***x***_*alt*_ in all position, and solve the resulting optimization problem using gradient descent. The final optimized *γ* will be used as positional effect score of the given motif.

#### Algorithm 1 Position Constrained Motif Generation

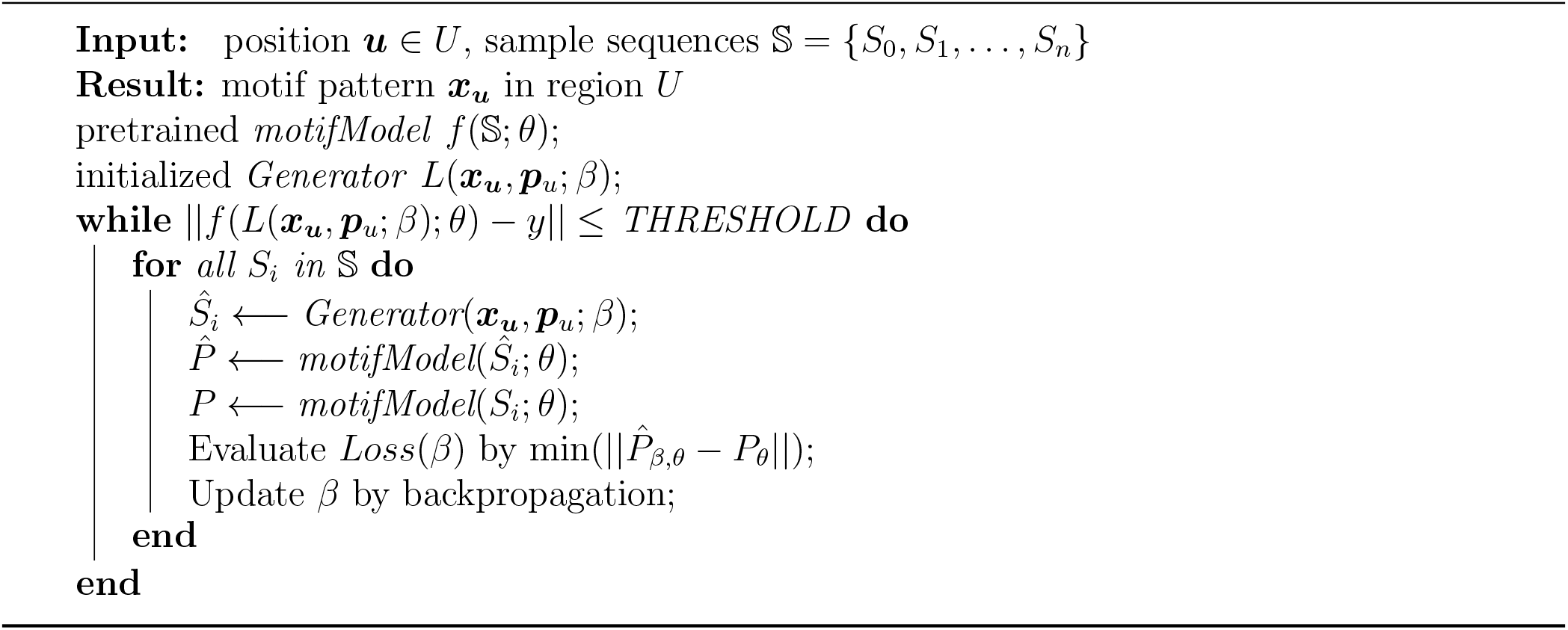

### 3.4 Motif generation constrained by position

If a region of certain position *U* in sequence is known as important, we can establish motif generation process as denoted in Algorithm (1) by minimizing the target loss of a pretrained model:

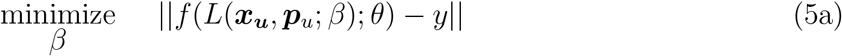

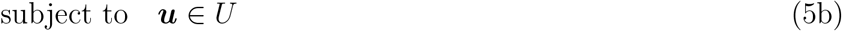

where *L* is a mapping function parameterized by *β* to update motif pattern for sequence input ***x***_***u***_. In practice, it is a generator network implicitly represent a sequence pattern by position ***p***_*u*_. When appropriately optimized, this generator can produce meaningful motif ***x***_***u***_ with the same distribution in ground true samples.

## 4 Experiments

Two model training experiments were conducted on mRNA translation and pre-mRNA splicing events tasks. In the first experiment, a translation model was trained on 280,000 synthesized mRNA sequences of 50 nucleotides in length from Sample et al. (2019). The translation efficiency values for these sequences were labeled based on cell line experiments. Then we evaluated model interpretation using variants in human 5’UTR mRNA sequences, which were downloaded from Lim et al. (2021). In the second experiment, a splicing model was trained on 20,287 pre-mRNA sequences from the GENCODE V24lift37 dataset released by Harrow et al. (2012). Followed the data prepossessing scheme in Jaganathan et al. (2019), we took splicing site annotations from canonical transcripts. The splicing probabilities for these sequences were labeled in the middle 5000 nucleotides.

Both models were trained in two stages. In the first stage, a base motif encoder was trained to predict the task label. The purpose of this stage was to obtain suitable parameters in *f* (***x***_*i*_; *θ*) for representing motif patterns related to the task. In the second stage, a positional encoder was trained by aligning activated motif features(denoted as motifs A) in the middle of the input sequence and applying a positional encoding function. Positional encoding function *sin*(***x***_*i*_; *γ*) was applied as suggested in Sitzmann et al. (2020); Vaswani et al. (2017). After the training of the second stage, the positional activated motifs (denoted as motifs B) were taken as the interacted sequence patterns.

### 4.1 Motifs pattern discovered in translation sequences

We discovered diverse motif from the optimized translation model. Some examples of these motifs are shown in Figure 3 and match common biological knowledge. For instance, we observed that upstream AUG/GUG(uAUG/uGUG) and UAA/UAG codon(motif B18 and A9) nearby in mRNA 5’UTR were interacted periodically in an interval of three. We can reason that co-occurring start and stop codons will form an upstream open reading frame, inducing mRNA to produce an extra short peptide. The order for these two motif is important. Only when a stop codon UAA/UAG in motif B18 occurs downstream of uAUG/uGUG will the translation efficiency be regulated. Further, GC-rich structural elements like sequence motif A26 can affect the function of uAUG (upstream AUG) in translation through their interaction with the mRNA sequence (Tang & Tseng, 1999). GC-rich regions are typically more stable than AT-rich regions, and can influence the secondary and tertiary structures of RNA molecules. In the case of uAUG, the presence of a GC-rich structural element nearby may affect the accessibility of the start codon to the ribosome, the molecular machine responsible for translating mRNA into protein. This can ultimately impact the efficiency of protein translation and the function of the resulting protein.

**Figure 3:**
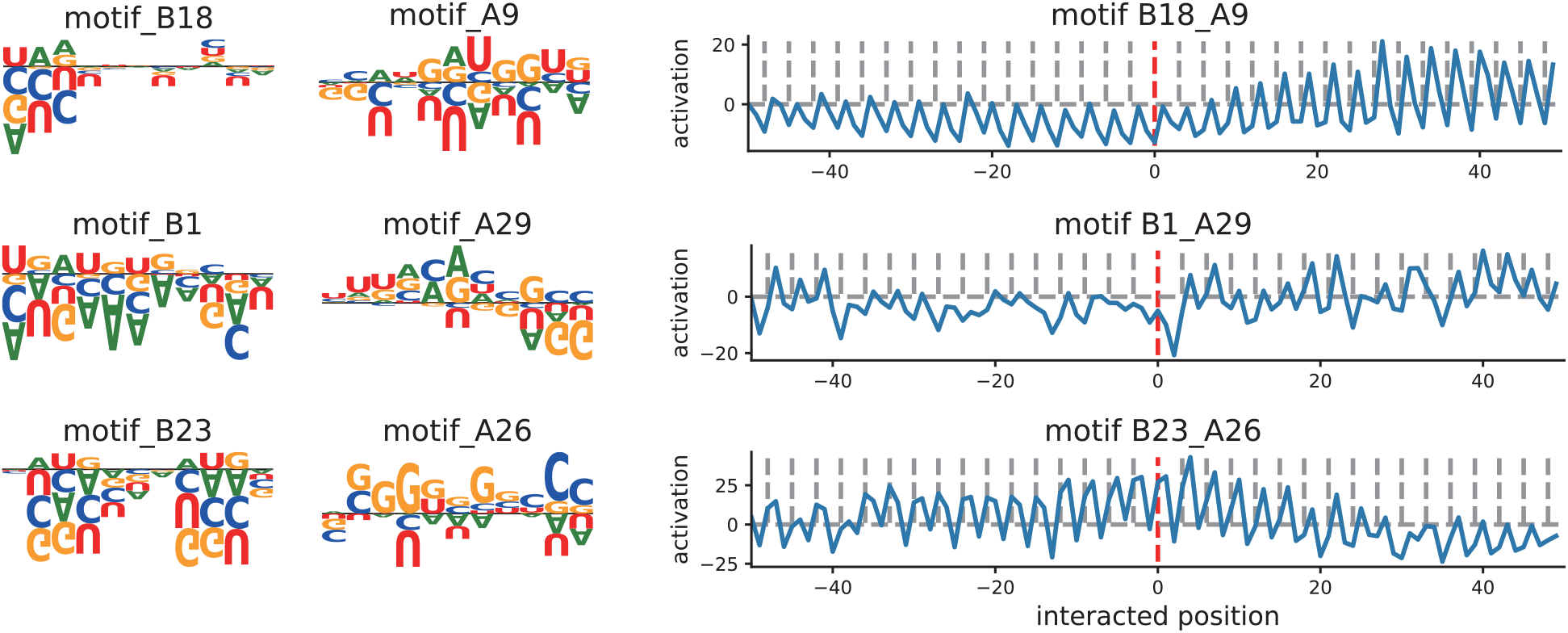
Specific examples taken from motifNet interaction matrix trained on 5’UTR mRNA data. A motifs were aligned in the middle(red line) and blue line plots represent the positional activation effect of B motifs.

We also noticed that the RBP motif GUAUGU, similar to the RNA sequence pattern binding to KHSRP from mCross database(Fig. 4b) by Feng et al. (2019), interacted with motif A29 [A/G]CCGCC. We suspected this fragment is related with GCC-box (sequence with GCCGCC pattern). It is reported by Prajapati et al. (2019) that GCC-box is an essential sequence for enhancement of gene expression.

**Figure 4:**
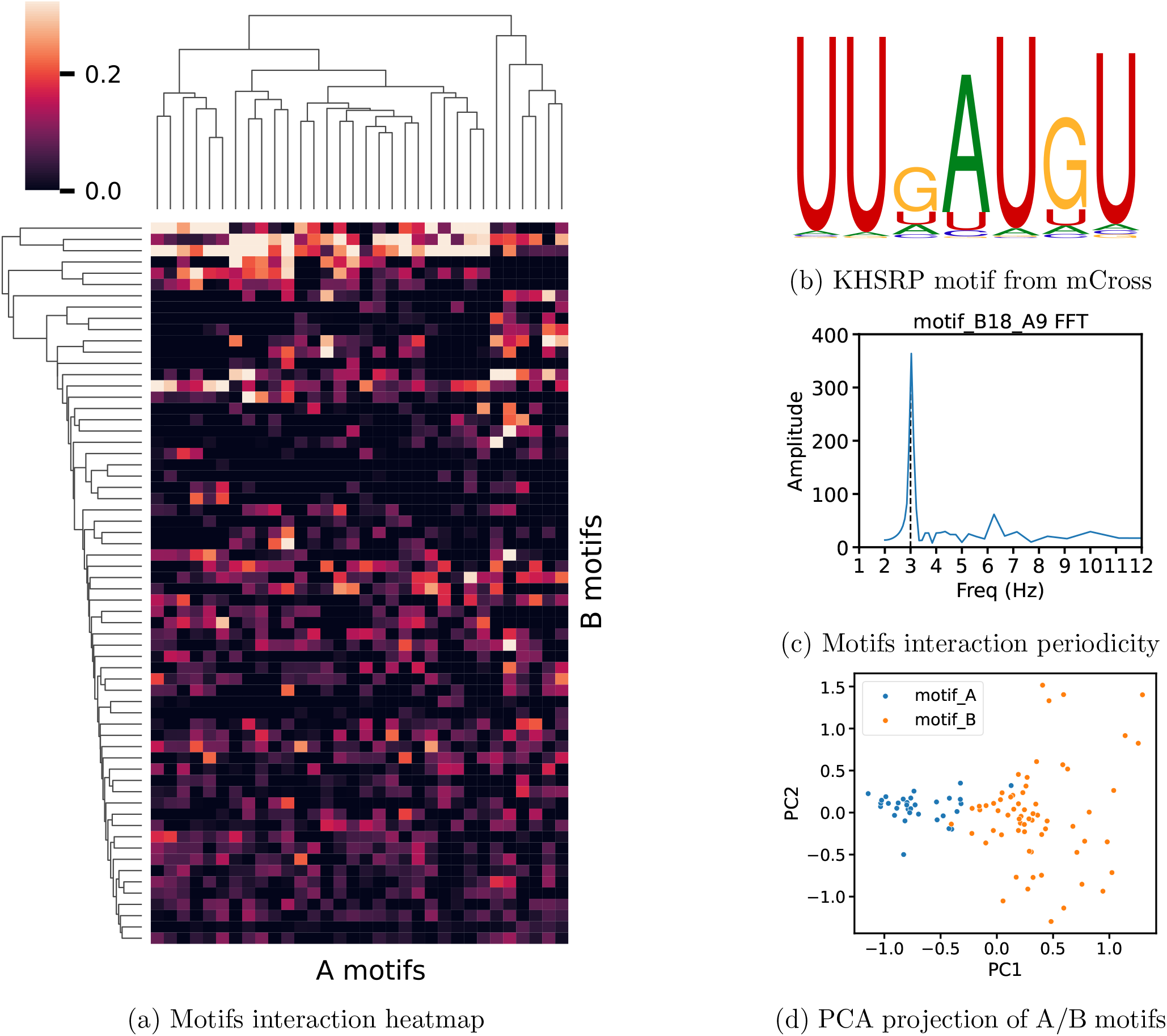
a) Visualization of pattern interaction matrix between A-B motifs. Patterns matched in positional activation were in brighten colors. b) KHSRP protein RNA binding motif selected from one of the top10 clustering patterns in mCross database. c) There is an obvious periodic interval at 3 in the frequency domain by Fourier transformation of positional activation. d) Patterns were significantly different between A/B motifs.

It was worth noticed that the above interacted pairs of motifs A and B were actually significantly distributed in two clusters (Fig. 4d). This indicated A motifs might directly influence final translation while B indirectly regulated translation efficiency by interaction with A, since our two stage training strategy produced A motifs by directly fitting target translation, and then produced co-interactive B motifs in the second training phrase.

Besides, we observed some variants in non-trivial positions that violate motif patterns resulting in negative translation downstream regulation. Examples shown in Fig. 6) demonstrate that most of the variants occurring in relative positions with high activation resulted in significant lower translation log fold change. We assumed that the paired motif elements possibly created new upstream open reading frames (uORFs), which are potent regulatory elements located in 5’UTR of mRNA transcript(Fernandes & Romão, 2020; Lin et al., 2019).

**Figure 5:**
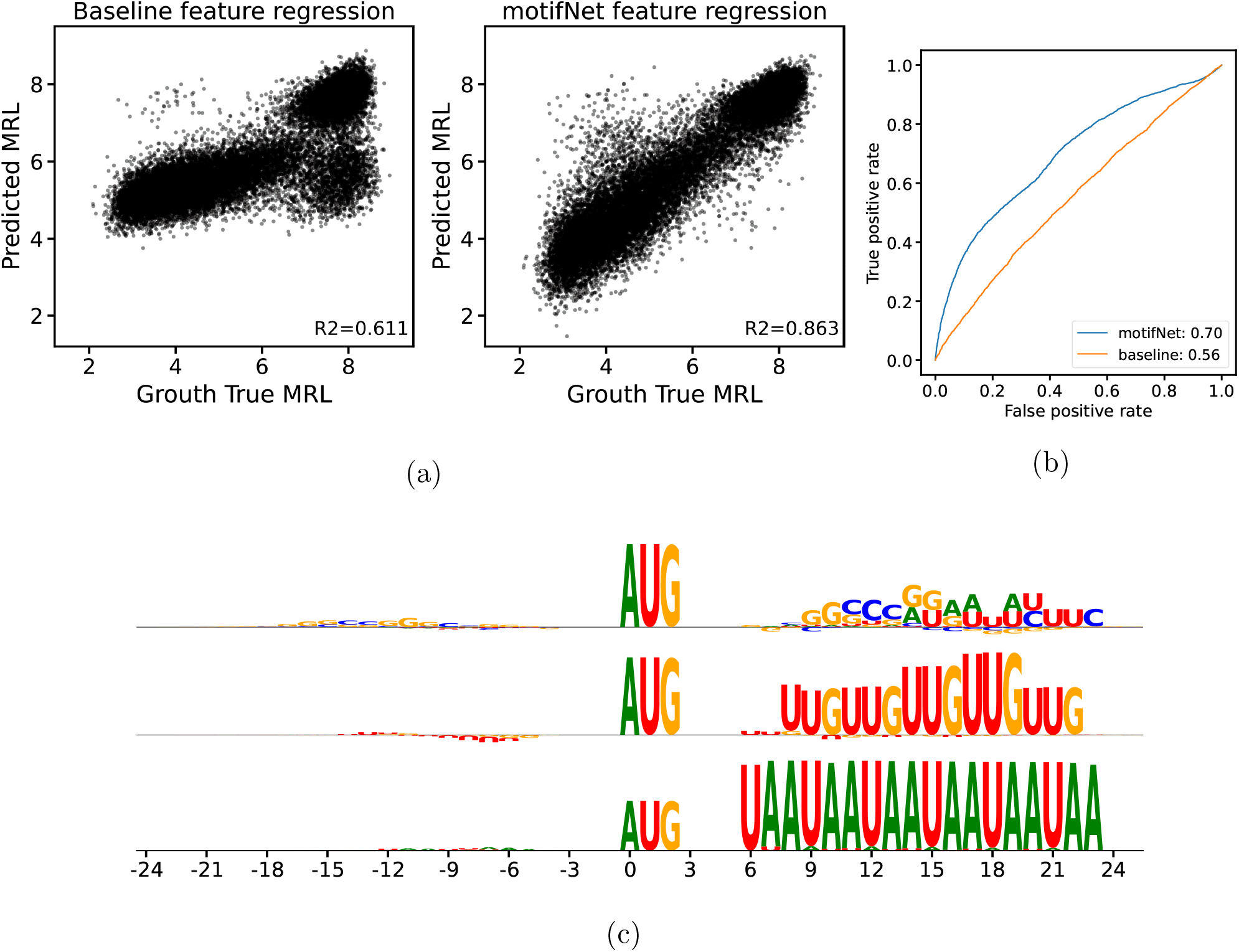
a) Performance of MRL prediction from linear regression model using motif feature. b) Accuracy and specificity ROC curve of a out-of-frame uAUG representation. c) Examples showing the activation of a out-of-frame uAUG motif was conditioned on interactions with other regulation elements. Letter were drawn according to the relative effect delta scores that altered the original uAUG. For example, GC-riched sequence were plotted in bigger size from relative position 9 to 15 since it dramatically decreased the motif activation score, resulted in a large score change. Letters upside down were shown to have the opposite effect.

**Figure 6:**
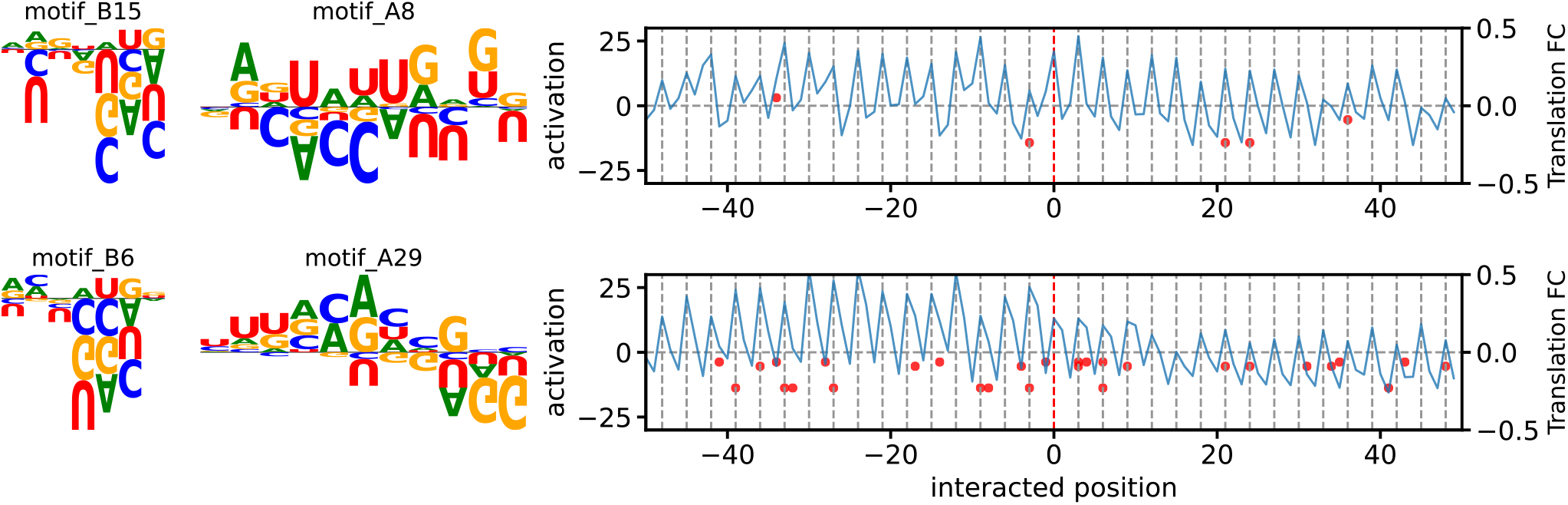
Position distribution of prostate cancer related single nucleotide variants(SNV) in human mRNA 5’UTR region. Blue line plots denoted the positional activation effect of motif B. Red dots represent pattern position(x-axis) and translation efficiency fold change(y-direction) caused by mutation in motif B in each SNV relative to motif A(red line).

### 4.2 Motifs representation evaluation and interpretation

To evaluate how well we predict the RNA motifs in translation model, we define positive examples as sequences that contain translation-regulating motifs and negative examples as sequences that do not contain these motifs or that contain motifs that have a different effect on translation. As the MRL prediction model can reach a performance with R-square greater than 0.9, we can use the model predicted translation value itself as labels. Take out-of-frame(OOF) AUG as an example, normally it will decrease translation level, as it form an wrong upstream ribosome initial site. In this scenery, this AUG motifs should be activated with high motif activation score as it have a great regulation on translation efficiency. However, it is also important to consider the potential impact of any out-of-frame AUG motifs can have different effects on translation efficiency depending on the context in which they occur. When an OOF AUG meets another stop codon, like UAA, it will form a complete upstream open reading frame to produce a short peptide. An extra short peptide was less translation regulated than a wrong translation initialization. So this can be treated as a negative sample, whose motif activation score should be low.

Therefore we measure the accuracy, precision of motifs representation using motif activation score versus mRNA event label. In this case, an absolute delta log2 fold change of translation efficiency greater than 1.0 caused by any random mutation to the motif was labeled as positive sample. In Figure 5 we noticed that motifNet activated motifs are more capable of considering incorporating additional features or information into account for context-dependent effects.

### 4.3 Motifs generation in splicing sequences

Splicing model is more difficult to train than translation model since the input sequences are much longer. SpliceAI model suggest using huge network structure for long term sequence information extraction. However, we found that our splice model kept a relatively good performance with much smaller network structure. Our validation top-k-accuracy was 85% by using only one layer of 1d convolution for motif encoding, with a 8-layers down sampling activation mask. We also experimentally trained a model using 3-6 layers as motif encoder, which increase the validation accuracy up to 87%. Considering the interpretation of motifs, we kept using one layer in motif encoder.

Given positional information as input, motifNet outputs sequence by optimizing the probability of splicing site in an implicit way. It is known that there are branch point sequence patterns within -40 to -20 in the upstream of an acceptor splicing site. We input this position to motifNet and there is an obvious branch point pattern UACUAAC occuring in the region (Fig 4.b). Furthermore, we also found that there is an AU-rich and AUGU pattern in this region, as previously discussed in 5’UTR part that this is a functional protein binding motif, such as KHSRP.

We feed the motif prior model with two inputs, TACTAAC branch point pattern and GC-rich sequence pattern. For the same branch point pattern, it was found to positively affect splicing among -40 to -20 relative positions in Jaganathan et al. (2019). In our experiment the results were similar, with the additional perspective that the TACTAAC pattern within -100 to -50 upstream will yield negative effects to splicing. Compared with Nazari et al. (2019) using both frequency distribution and deep learning model, our motifNet application in splicing performed much better with more motif patterns.

We also studied the GC-rich pattern effect on splicing since it is reported in Georgakopoulos-Soares et al. (2019) that G4 and GC-rich patterns act as splicing modulator in exon region. We took an intuitive pattern GGGGCCCGGGG as example, the positional score indicates that it will produce an extreme negative effect but among -100 to -50 upstream regions, it will help promote the splicing. This might possibly be due to the mechanism that stable structure enhances pre-mRNA stability for splicing regulatory protein binding. We further investigate one of the G4 elements, (*GGGNNN*)_3_*GGG* as splicing modulator in exon region, as reported by (Georgakopoulos-Soares et al., 2019), according to Georgakopoulos-Soares et al. (2019), 35% of human genes contain a G4-seq peak within 100 bp of a splice junction, supporting our observations in Fig. 8.

**Figure 7:**
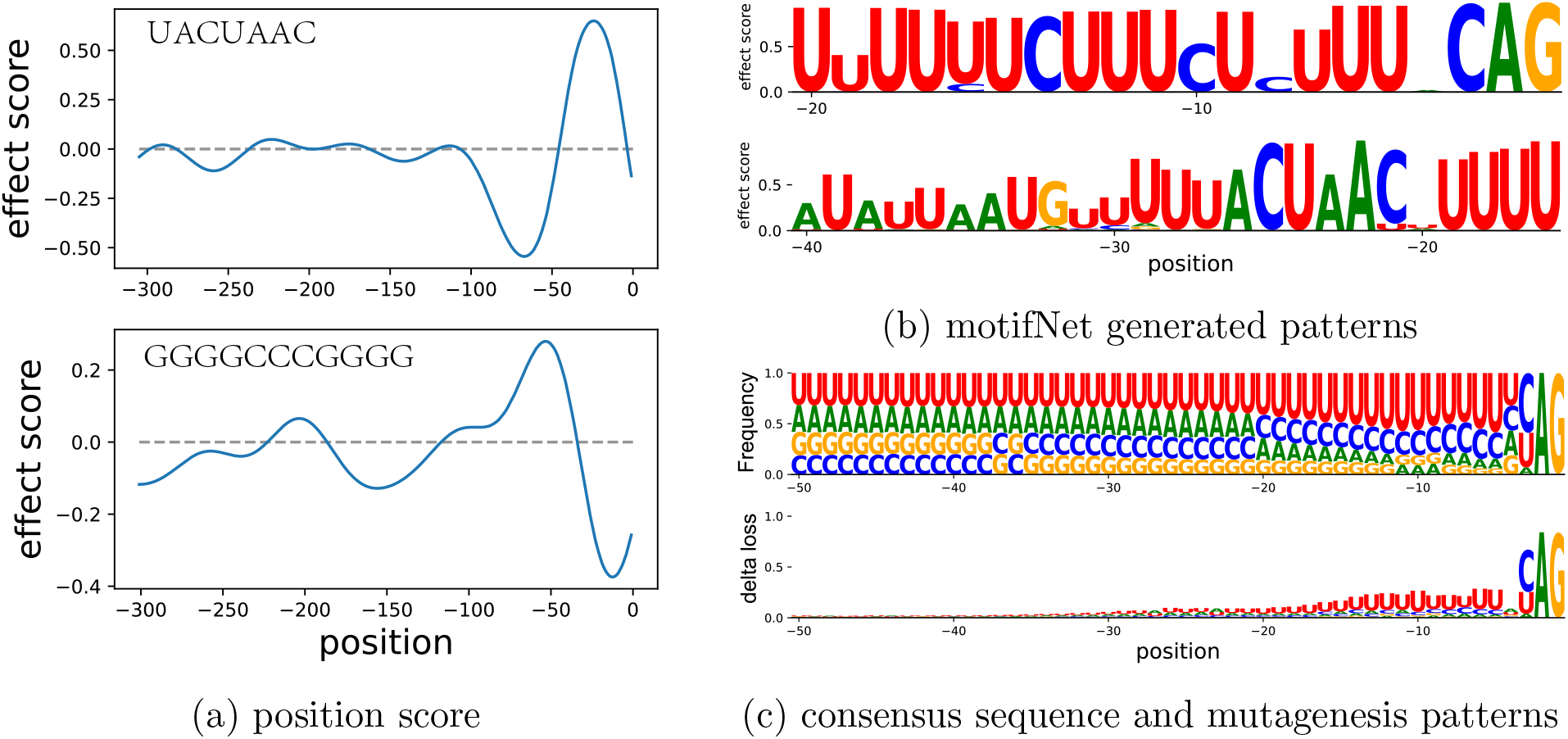
Sequence patterns discovered in implicit splicing model. a) Effect score generation condition on branch point pattern UACUAAC and GC-rich motif. b) Motif generation constrained on position c) Motif patterns extraction using frequency statistic based on all aligned acceptor sequences in training dataset and single nucleotide mutation over the whole upstream acceptor splicing site(mutagensis) using spliceAI as benchmark.

**Figure 8:**
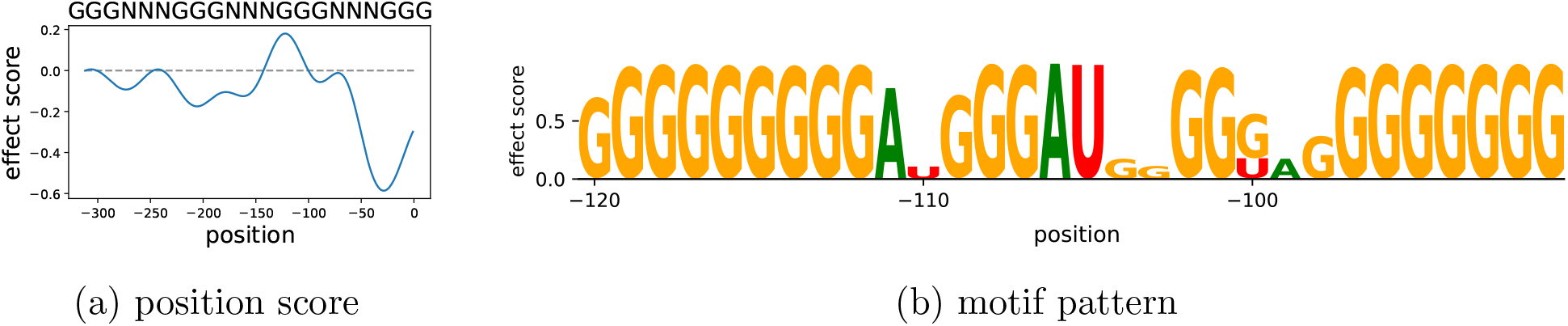
G4 pattern validation in motifNet splicing model. a)Sequence wide effect score generation constrained on G4 motif. b) Motif generation constrained on coordinate between -120 to -90 region relative to the acceptor splicing site.

## 5 Discussion

In summary, motifNet is specifically designed for mRNA motif interaction discovery using implicit neural representations. We defined the general framework where sequence patterns and coordinate information were separately modeled and solved. For traditional motif analysis approaches, researchers need to use domain knowledge to analyze frequency enrichment from sample sequences. As for motifNet, we are capable of generating motif interaction patterns constrained on positional inputs without predefined rules. We prototype models with translation and splicing labeled sequences and demonstrate examples of interpretative motifs matched with known reported biological patterns. Beyond classical mechanisms, we also explore unknown sequence variants potentially affecting human disease to validate the generalization of patterns. There will be promising applications in mRNA therapeutics(Kitamura & Nimura, 2021) if future studies can further extend this approach to identify more undiscovered driver mutations in the human genome.

## Part I Appendix

### A Model details

#### A.1 training

Here we provide some more details on the models that we use. For all sub modules in motifNet we use the Adam optimizer and learning rate was set to 0.001 for motif encoder, 0.0002 for splicing motif/positional activation generation. Batch size of training is 32 for splicing dataset and 128 for translation dataset.

Activated motifs number is hyper-parameter defined by users. Here for translation model we set 32/64 for A/B motifs, and for splicing model we set 32/32 for A/B motifs.

Motifs/position constrained generation was only applied in splicing data since this can only be trained from real human genome annotation datasets. Translation model was not applicable on this task since training sequences from Sample et al. (2019) were random synthesized. Given better datasets with human translation labels in future studies, same task can be applied in this field.

#### A.2 Environment

- Torch version: 1.10.0
- Cuda release: 11.5, V11.5.50

Other environment variables not affecting model results are listed here.

Models are all trained in a single GPU. The maximum training time is no than 12 hours.

#### A.3 model flowchart

**Figure 9:**
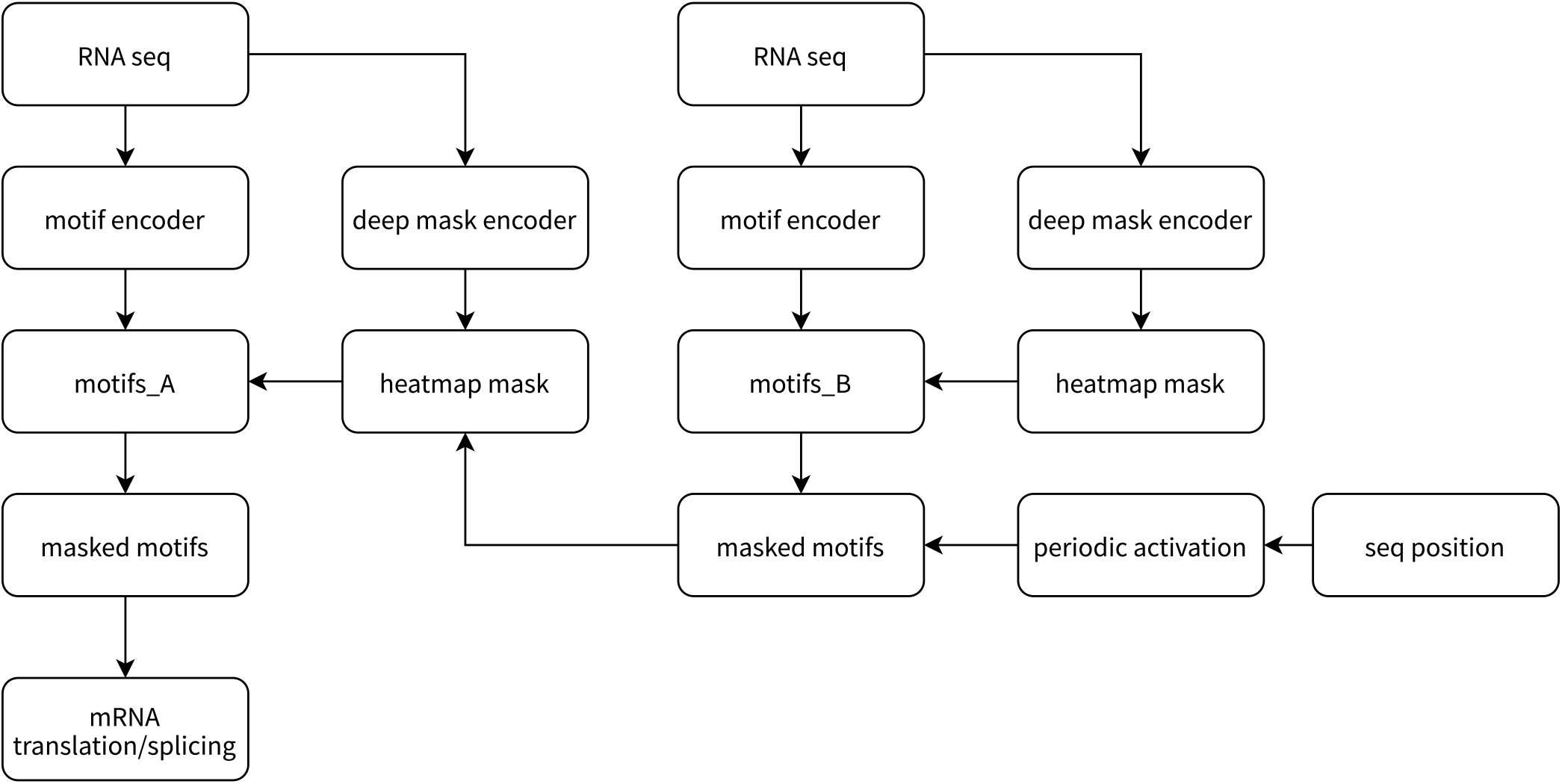
Two stage motif fitting network.

### B Motif patterns in translation model

#### B.1 Motifs interaction visualization

**Figure 10:**
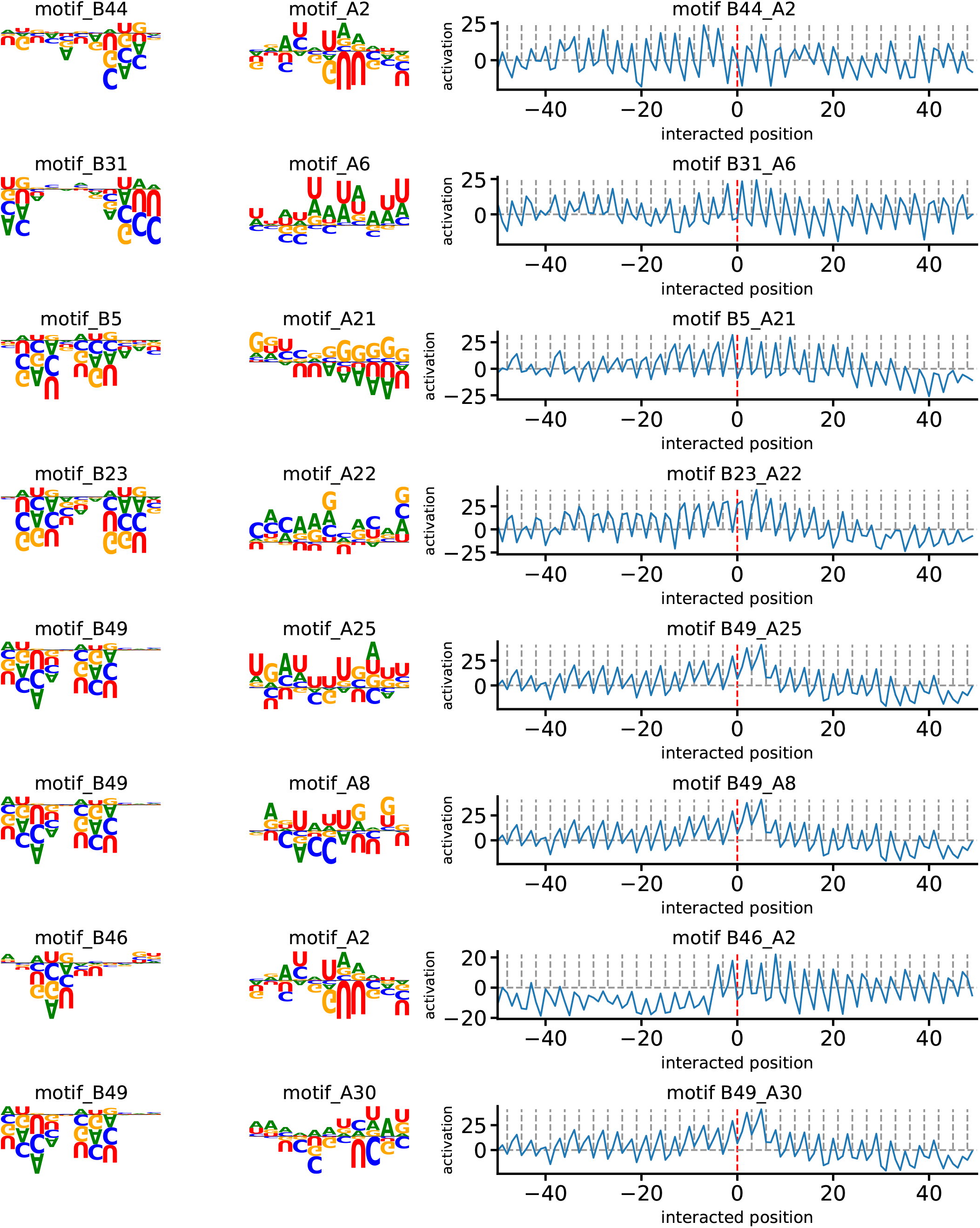
Motifs interacted pairs with high activation score.

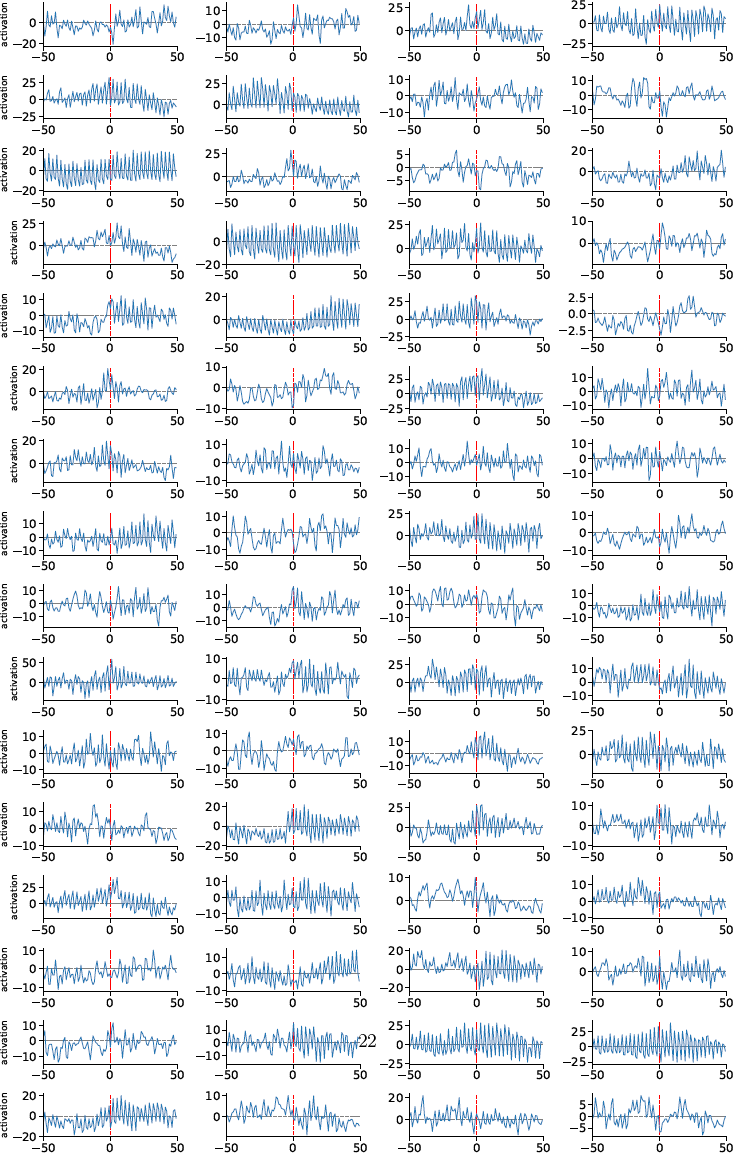
All interaction position activation of 5’UTR motifs

### C Motif patterns in splicing model

**Figure 12:**
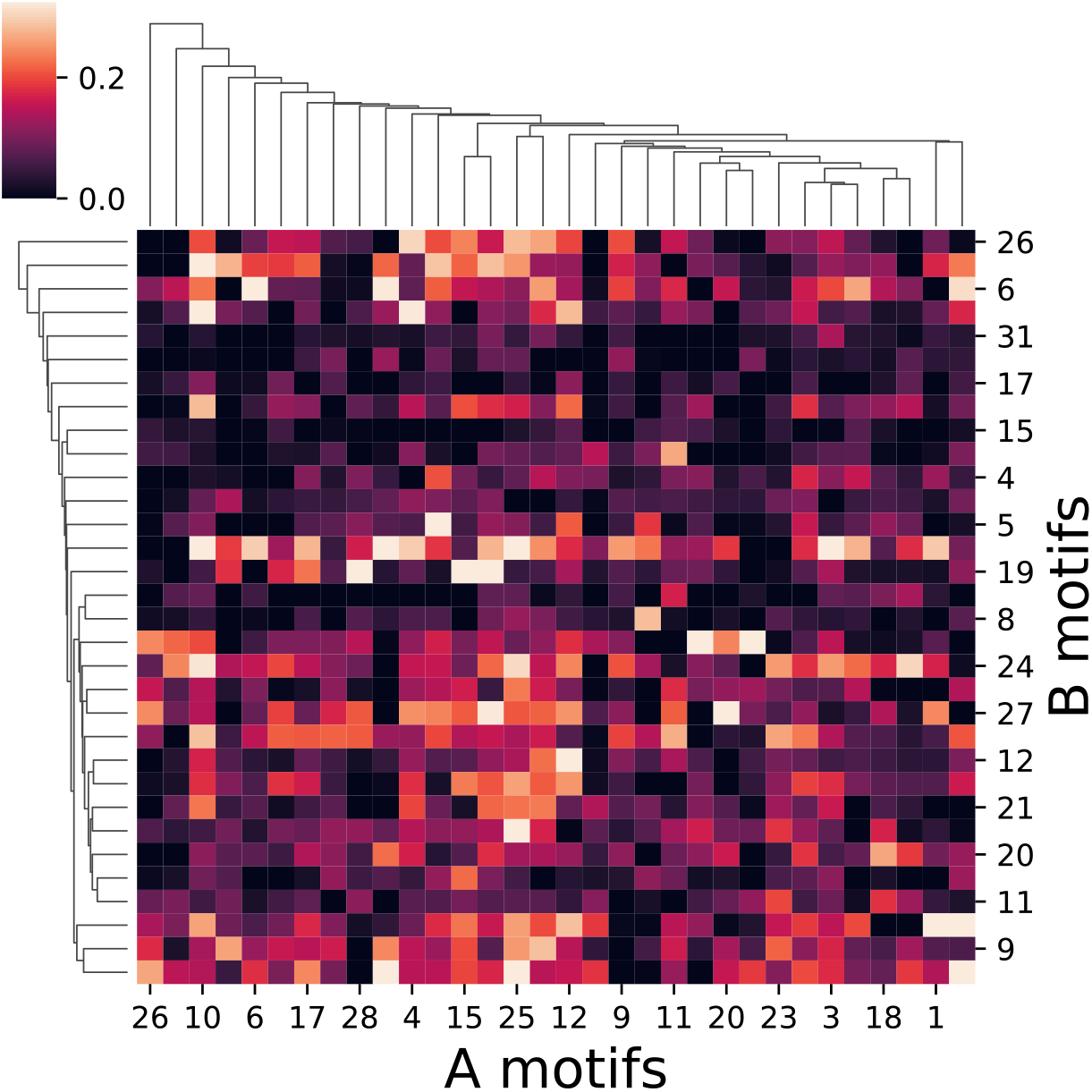
Visualization of pattern interaction matrix between A-B motifs. Patterns matched in positional activation were in brighten colors.

**Figure 13:**
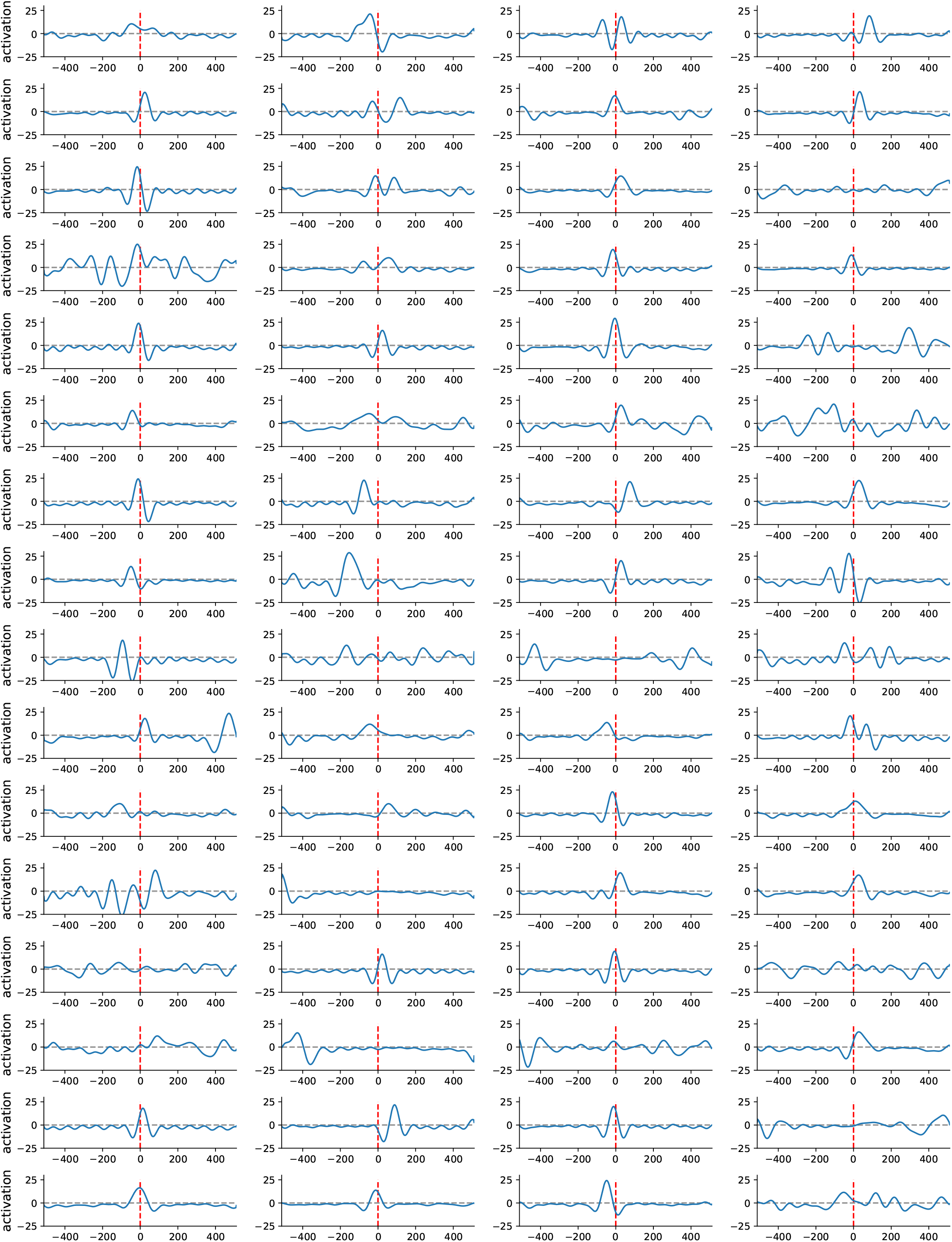
All interaction position activation of splicing motifs 24

